# Median eminence myelin continuously turns over in adult mice

**DOI:** 10.1101/2022.11.16.516753

**Authors:** Sophie Buller, Sara Kohnke, Robert Hansford, Takahiro Shimizu, William D Richardson, Clemence Blouet

**Author notes:** The authors have no interest to disclose.

## Abstract

**Objective:** Oligodendrocyte progenitor cell differentiation is regulated by nutritional signals in the adult median eminence (ME), but the consequences on local myelination are unknown. The aim of this study was to characterise myelin plasticity in the ME of adult mice in health or in response to chronic nutritional challenge.

**Methods:** We assessed new oligodendrocyte and myelin generation and stability in the ME of healthy adult male mice using bromodeoxyuridine labelling and genetic fate mapping tools. We assessed the contribution of microglia to ME myelin plasticity in PLX5622-treated C57BL6/J mice and in *Pdgfra-Cre/ER^T2^;R26R-eYFP;Myrf^fl/fl^* mice, where adult oligodendrogenesis is genetically blunted. Finally, we investigated how 45% high fat diet or 70% caloric restriction feeding paradigms impact ME oligodendrocyte lineage progression and myelination in C57BL6/J mice.

**Results:** We show that myelinating oligodendrocytes (OLs) are continuously and rapidly generated in the adult ME. Paradoxically, OL number and myelin amounts remain remarkably stable in the adult ME. In fact, the high rate of new OL and myelin generation in the ME is offset by continuous turnover of both. We show that microglia are required for continuous OL and myelin production, and that ME myelin plasticity regulates the recruitment of local immune cells. Finally, we provide evidence that ME myelination is regulated by the body’s energetic status, decreased in calorie-restricted animals, and increased in mice fed a high fat diet.

**Conclusions:** This study uncovers a previously unappreciated form of ME structural plasticity and mechanism of myelin remodeling in the adult brain.

## 1. INTRODUCTION

Oligodendrocytes (OLs) are the myelin forming cells of the central nervous system (CNS). Once thought to be exclusively generated during early postnatal life in rodents, it is now evident that new myelinating OLs continue to differentiate from a pool of oligodendrocyte progenitor cells (OPCs) in various white matter tracts of the adult brain long after developmental myelination is complete (*1–6*). In addition to facilitating brain repair after injury (*7*), new OL and myelin generation in the adult brain optimizes physiological processes including motor skill learning (*8, 9*) and memory processing (*10, 11*). Previous studies have demonstrated that new OLs generated during adulthood are long-lived and stably integrate into existing circuits (*12*) where they either contribute towards the modification of existing myelinated circuits, termed myelin remodeling, or myelinate previously naked axons, termed de novo myelination (*13–15*). However, the rate of OPC proliferation and differentiation, and therefore new OL generation, varies by brain region with, for example, adult oligodendrogenesis occurring at a greater rate in the corpus callosum (CC) than motor cortex (*2*).

The median eminence (ME) is a circumventricular organ located adjacent to the arcuate nucleus (ARC) at the base of the hypothalamus, a region critical for homeostatic functions. Due to its unique fenestrated vasculature, the ME maintains open communication with the circulation, allowing the release of hypophysiotropic hormones directly into the portal system, and unbuffered exposure to circulating factors for optimal monitoring of circulating cues (Rodriguez et al., 2010). The ME is highly plastic, undergoing cellular and structural remodelling that is critical for adaptive physiological responses to stimuli such as energy deficit (*17*) and the regulation of the neuroendocrine axes (*18*). Although the hypothalamus is largely devoid of myelin, myelinating OLs are present in the adult ME. Here, myelin can be observed in a dense band that extends laterally across the dorsal ME, directly below the tanycytes lining the wall of the third ventricle, where it ensheathes axons of magnocellular neurones (*19, 20*).

In the ME, OPCs proliferate more rapidly than in adjacent hypothalamic nuclei *in vivo* (*6, 21*), and the proportion of differentiating OPCs is higher and regulated by nutritional signals (*19*). However, the consequences on local myelination are unknown. The initial goal of this study was to quantify adult oligodendrogenesis in the ME OL compared to the CC, a well-characterised white matter tract where continuous OL generation occurs in adulthood (*2*). Using bromodeoxyuridine (BrdU) and genetic fate mapping approaches, we demonstrate that new myelinating OLs are rapidly and continuously produced in the healthy adult ME, and that new OL and myelin generation are offset by the continuous turnover of myelinating OLs which maintains stable OL and myelin density in the ME over time. Mechanistically, we show that microglia are essential for the maintenance of ME OL and myelin production and that OL plasticity is required for the maintenance of the local immune cell populations. Finally, through the assessment of myelination and OL lineage plasticity in the ME of mice exposed to a high fat diet (HFD) or caloric restriction (CR), we provide evidence that ME myelin amounts are regulated by peripheral energy availability.

## 2. MATERIALS AND METHODS

**Resource table**

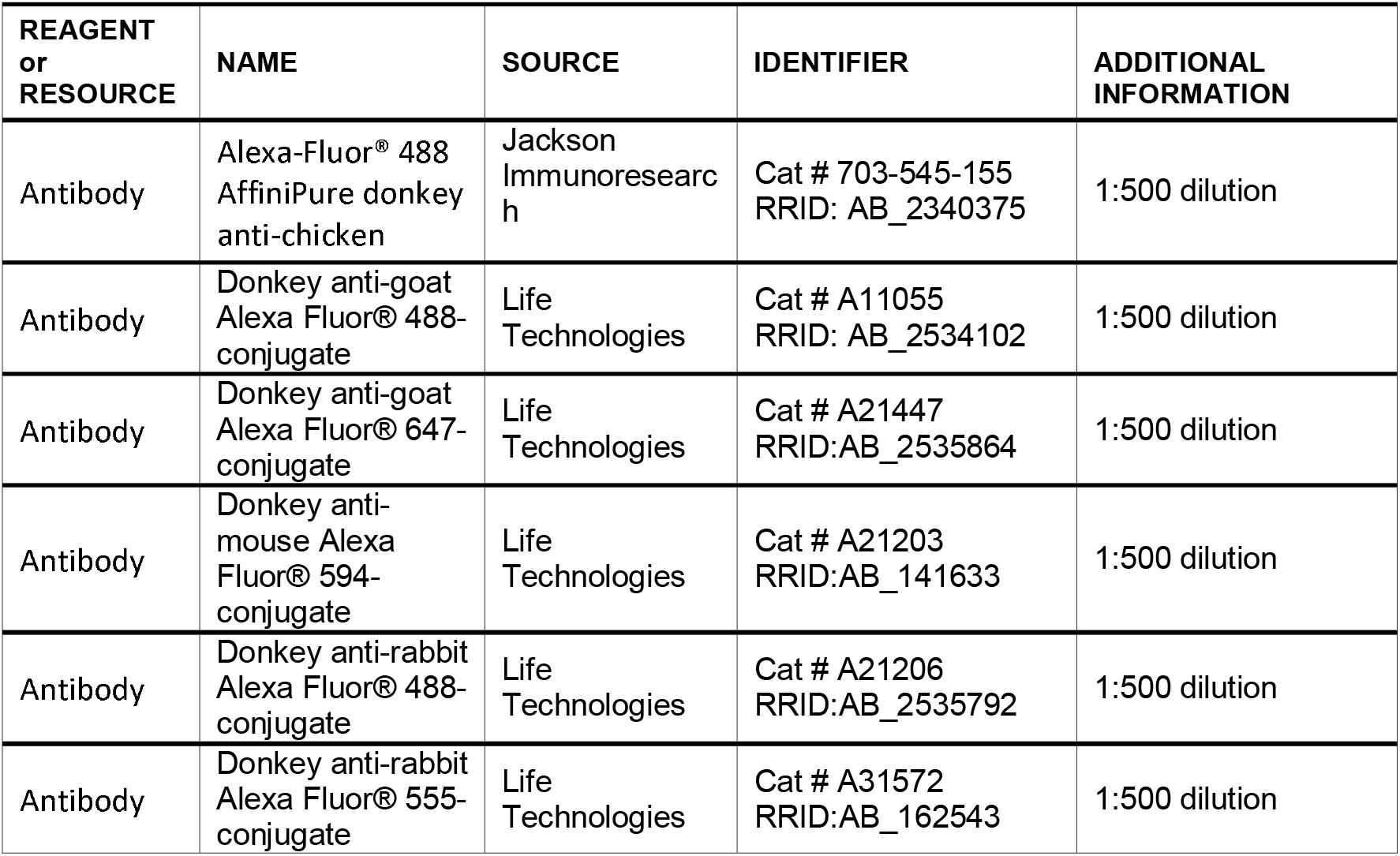
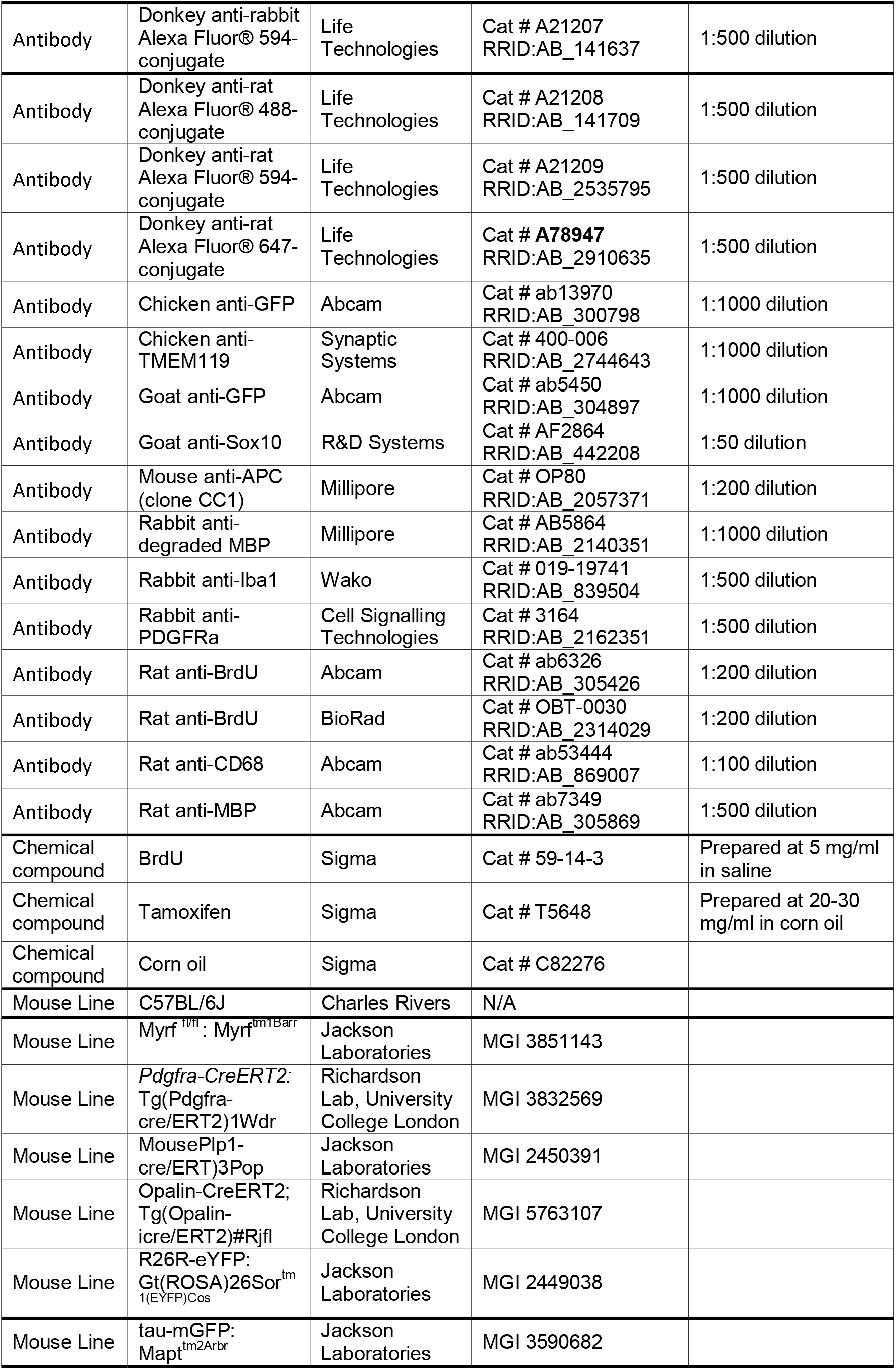
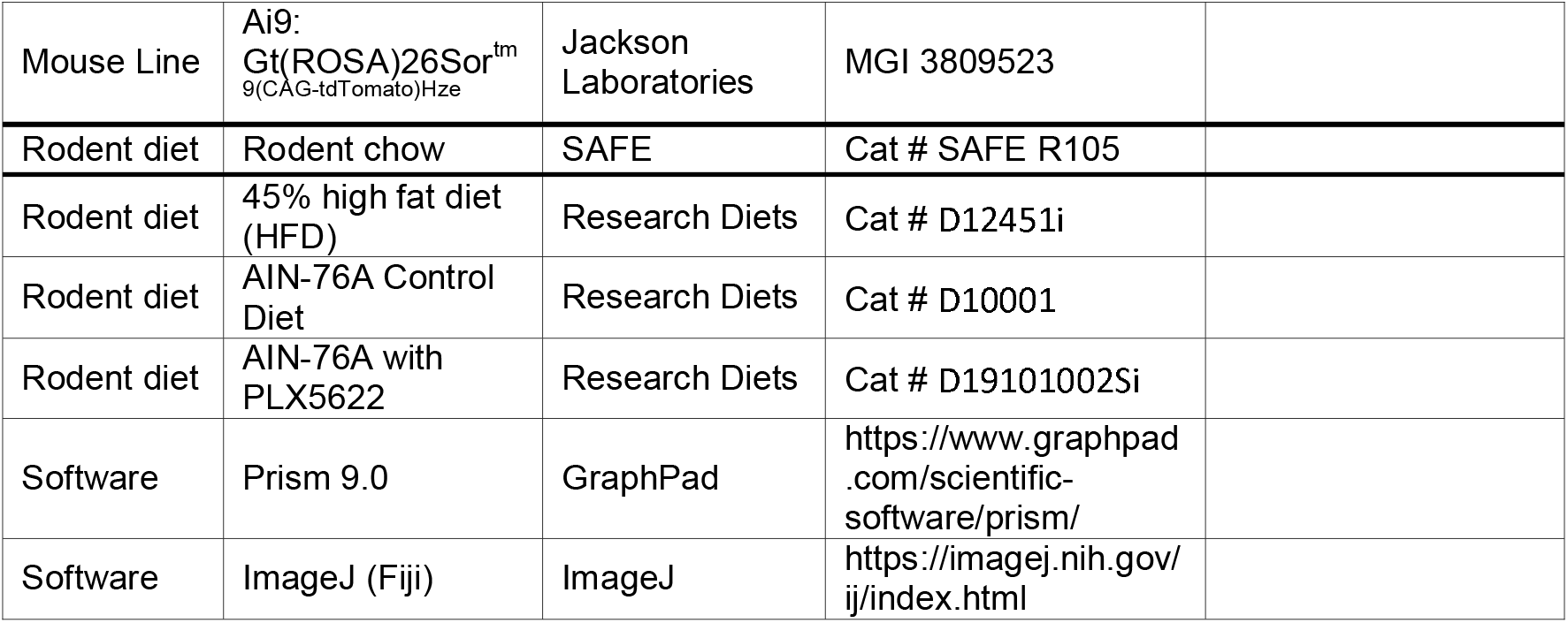

### 2.1. Animals

All animal experiments were performed in accordance with the UK Home Office regulations under the Animals (Scientific Procedures) Act (1986) and with the approval of the University of Cambridge Animal Welfare and Ethics Review Board. Animals were group-housed in a specific pathogen free facility and maintained on a standard 12-hour light/dark cycle (lights on 7:00-19:00) at 22°C with *ad libitum* access to water and standard laboratory chow (SAFE R105, SAFE Complete Care Competence, Rosenberg, Germany) unless otherwise stated. All experiments were performed on male mice starting from postnatal day 60 (P60). C57BL/6J mice were obtained from Charles Rivers Laboratories (Saffron Walden, UK). *Pdgfrα-Cre/ER^T2^;Rosa26-YFP;Myrf^fl/fl^* mice were provided by William Richardson at University College London. *Opalin-Cre/ER^T2^;Ai9* mice were provided by Professor Thora Karadottir at the University of Cambridge. *Plp-Cre/ER^T2^* mice (stock number 005975) and B6;129P2-*Mapt*^*tm2Arbr*^ /J (*tau-mGFP*) mice (stock number 021162) were obtained from the Jackson Laboratories (Bar Harbour, Maine).

### 2.2. Tamoxifen Preparation and Administration

Tamoxifen (Sigma, St Louis, Missouri) was prepared in corn oil (Sigma) by sonication at 37°C at 30 mg/ml prior to administration by oral gavage at 300 mg/kg on 4 consecutive days. Where tamoxifen was administered by the intraperitoneal route, tamoxifen was prepared at 20 mg/ml for injection at 80 mg/kg for 8 consecutive days.

### 2.3. Cumulative bromodeoxyuridine (BrdU) paradigm

Mice were administered BrdU (Sigma), dissolved in drinking water at 1 mg/ml continuously for up to 60 hours and concomitantly administered up to two intraperitoneal injections of BrdU (50 mg/kg; prepared in sterile saline at 5 mg/ml) in any 12-hour period. Mice were sacrificed 2, 12, 24 and 60 hours after the onset of BrdU administration.

### 2.4. Caloric-restriction paradigm

C57BL/6J mice were acclimatised to single-housing and randomly allocated to the ad libitum fed or calorie restricted group. For animals that were calorie-restricted, 70% of the average amount of food eaten by the control group over the previous 24-hour period was provided at ZT11 daily. 7 days after the onset of calorie restriction, all animals were administered four intraperitoneal injections of BrdU (50 mg/kg) within a 24-hour period starting at ZT5 and ending at ZT4 on the eighth experimental day. Mice were sacrificed one hour later, at ZT5, on experimental day 8.

### 2.5. 45% High Fat Diet Studies

C57BL/6J mice were fed a diet with 45% of calories from fat (45% HFD - D12451, Research Diets, New Brunswick, New Jersey) or a standard chow diet for 8 weeks from 7-8 weeks of age. Prior to culling, animals were administered 4 intraperitoneal injections of BrdU (50 mg/kg) over 24 hours as described above.

For fate mapping studies in *Plp-Cre/ER*^*T2*^*;R26R-eGFP* mice, animals were fed a 45% HFD from P60 and were administered tamoxifen 1-or 8-weeks later. Mice were trans-cardially perfused 14 days after tamoxifen administration.

### 2.6. Microglia Ablation with PLX5622

C57BL/6J mice were fed a control AIN-76A diet (D10001, Research Diets) or AIN-76A with PLX5622 at 12,000 parts per million (D19101002Si, Research Diets) from 8 weeks of age for 2 weeks prior to perfusion fixation.

### 2.7. Perfusion Fixation

Mice were administered 50 ul pentobarbital (Dolethal, 200 mg/ml) by intraperitoneal injection to achieve deep anaesthesia. Mice were then trans-cardially perfused with 50 ml heparinised phosphate buffered saline (PBS) followed by 4% paraformaldehyde (PFA; Fisher Scientific, Waltham, Massachusetts) in PBS (pH 7.4) at a flow rate of 5 ml/min.

### 2.8. Immunofluorescence

Brains were post-fixed overnight at 4’C in 4% PFA then cryoprotected in 30% (w/v) sucrose (Fisher Scientific) solution in PBS for at least 48 hours prior to processing. Tissues were covered with Optimal Cutting Temperature (OCT) medium (CellPath, Newtown, UK) and sections were obtained at 30 μm on a Leica SM2010R Freezing Microtome (Leica, Wetzlar, Germany). All sections were subjected to heat-mediated antigen retrieval in 10 mM sodium citrate (pH6.5; Fisher Scientific) in distilled water for 20 minutes at 80°C prior to washing 3 times in PBS. For sections immunolabelled for BrdU, tissues were then incubated with 2N hydrochloric acid (Sigma) in distilled water for 30 minutes at 37°C. Sections were subsequently incubated with 0.1M sodium borate (pH 8.5; Sigma) in distilled water to neutralise the acid and then washed 3 times with PBS. For all experiments, sections were blocked in normal donkey serum (NDS, Vector Biolabs, Philadelphia, Pennsylvania) diluted in PBS containing 0.3% Triton X-100 (0.3% PBST; Sigma) for one hour prior to primary antibody incubation overnight at 4°C. Primary antibodies were mouse anti-APC (OP80, Millipore, Burlington, Massachusetts) 1:500, rat anti-BrdU (ab6326 Abcam, Cambridge, UK) 1:200, rat anti-CD68 (ab53444 Abcam) 1:100, rabbit anti-degraded MBP (AB5864 Millipore) 1:1000, chicken anti-GFP (ab13970 Abcam) 1:1000, rabbit anti-Iba1 (Wako 019-19741, Richmond, Virginia) 1:500, rat anti-MBP (ab7349 Abcam) 1:500, rabbit anti-PDGFRα (3164 Cell Signalling Technologies, Danvers, Massachusetts) 1:500 and goat anti-Sox10 (AF2864 R&D Systems, Minneapolis, Minnesota) 1:50. Following primary antibody incubation, sections were washed 3 time with 0.1% PBST and incubated with appropriate fluorophore-conjugated secondary antibodies diluted 1:500 in 0.3% PBST for 2 hours at room temperature. Sections were subsequently washed with 0.1% PBST and mounted to Clarity microscope slides (Dixon Science, Edenbridge, UK) under coverslips (1.0 thickness; Marienfeld, Lauda-Königshofen, Germany) with Vectashield Vibrance Mounting Medium with 4′,6-diamidino-2-phenylindole (DAPI; Vector Laboratories, Newark, California). Alexa405, Alexa488, Alexa555, Alexa594 and Alexa647 conjugates (Life Technologies, Carlsbad, California) were used as secondary antibodies.

### 2.9. Confocal microscopy and image analysis

For all experiments, slides and images were blinded to experimental condition. Sections were imaged with a Leica SP8 confocal microscope using either a 40x or 63x oil objective. Sections were imaged as z-stacks at intervals of 3.3 μm with tile scanning to obtain signal from the entire depth and area of the region of interest (ROI). Microscope settings were identical for image acquisition within each experiment. Images were analysed using Fiji software. For all analyses, Z stacks were projected into a single image and the area of ROIs were selected using the freehand tool and measured. For cell counts, the Fiji manual cell counter was used to count marker-positive cells. Where area of immunoreactivity was calculated, all images within an experiment were identically thresholded and the area fraction, limited to that threshold, measured. 3D reconstruction images were generated using Imaris software.

### 2.10. Oil Red O staining

PFA-fixed tissues were cryoprotected in 30% sucrose solution (w/v in distilled water; Sigma) overnight before sectioning at 8-10 μm on a Leica CM1950 cryostat onto electrostatically charged slides. A stock solution of ORO was made up by gently heating 0.5 g ORO (Sigma) in 100 ml absolute isopropyl alcohol (Sigma) in a water bath overnight. A working solution was prepared by mixing 60 ml ORO stock solution and 40 ml 1% dextrin (w/v; Fisher Scientific) in distilled water. The working solution was allowed to stand for one day and filtered prior to use. Slides were rinsed in PBS and incubated with the ORO working solution for 20 minutes. Excess stain was rinsed off with distilled water and sections counterstained with haematoxylin for 20 seconds and blued in tap water. Coverslips were mounted with Pertex mounting medium (Pioneer Research Chemicals Ltd.).

### 2.11. Brightfield microscopy

Images were acquired with an Axioscan Z1 Slide Scanner (Zeiss) using a 20x objective.

### 2.12. Statistical analysis

All data visualisation and statistical analysis was performed in Prism 9 Software (GraphPad). Details of statistical tests are found in figure legends. All data are presented as mean ± standard error of the mean (S.E.M).

## 3. RESULTS

### 3.1. New oligodendrocytes are rapidly and continuously produced in the adult median eminence

We first characterised the cell cycle dynamics of OPCs *in vivo*, using a BrdU pulse-chase paradigm and immunostaining against pan-OL marker SRY-box transcription factor 10 (Sox10) and OPC marker platelet derived growth factor receptor alpha (PDGFRα) to visualise proliferative OPCs (Sox10^+^/PDGFRα^+^/BrdU^+^) in the ME, ARC and CC at postnatal day 60 (P60). In line with previous reports of rapid BrdU incorporation into ME OPCs (*6,21*), ME OPC total cell cycle time (T_C_) was ∼3.4 days, two to three times shorter than OPC T_c_ in the CC or ARC, (**Supplementary Figure 1A-B**), indicating rapid proliferation of ME OPCs. Unlike ARC and CC OPCs, ME BrdU^+^ OPCs rapidly lost PDGFRα expression (**Supplementary Figure 1C**) and gained expression of post-mitotic OL marker adenomatous polyposis coli (APC - also known as CC1) following 24-or 60-hours BrdU administration (**Figure 1A-B**), suggesting that adult-born OLs are rapidly and continuously produced in the ME.

**Figure 1.**
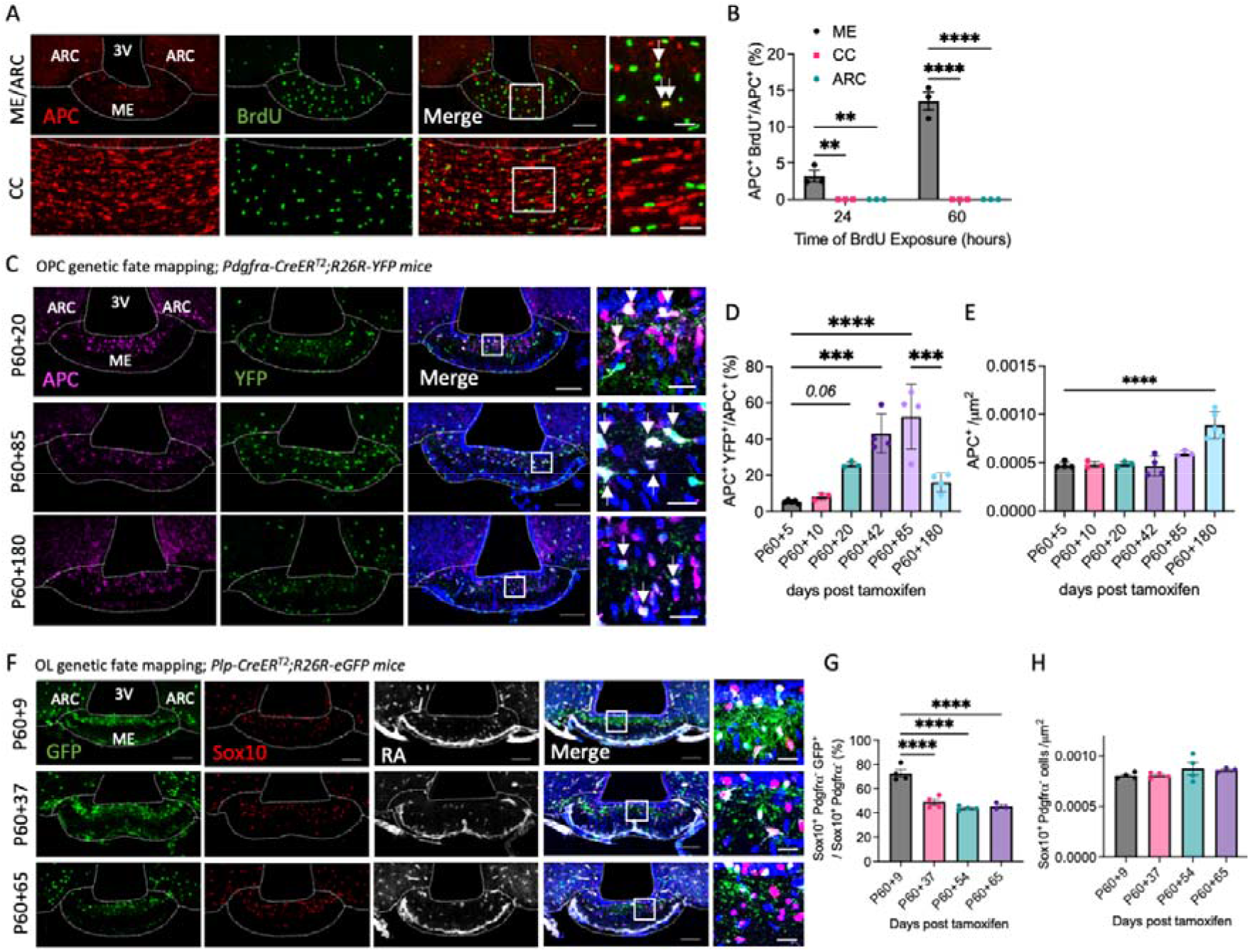
Rapid generation and turnover of oligodendrocytes in the healthy adult median eminence. Representative images of OL marker and BrdU or YFP/GFP expression in the ME of (**A**) 60 hour BrdU treated C57BL/6J -, (**C**) *Pdgfra-Cre/ER*^*T2*^*;R26R-YFP –* and (**F**) *Plp-Cre/ER*^*T2*^*;R26R-eG*FP-mice and associated quantifications (**B, D, E, G, H**). For all images, overview scale bars = 100 μm, inset scale bars = 20 μm. Arrows indicate adult-born OLs. Data (mean ± SEM) were analysed by one-way ANOVA with Dunnett’s multiple comparisons test or two-way ANOVA with Sidak’s multiple comparisons test, **p<0.01, ***p<0.001, ****p<0.0001, n=3-4/group.

To further establish rapid oligodendrogenesis in the adult ME *in vivo*, we labelled adult OPCs using *Pdgfrα-Cre/ER*^*T2*^*;Rosa26-YFP* mice administered tamoxifen at P60 and followed their fate over time by immunolabelling against OL markers and yellow fluorescent protein (YFP). Consistent with results from BrdU incorporation studies, labelled OPCs (Sox10^+^/PDGFRa^+^/YFP^+^) lost PDGFRa expression (**Supplementary Figure 1D-E**) and gained APC expression faster in the ME than in the CC, indicating rapid new OL production. Strikingly, adult-born OLs (APC^+^/YFP^+^) represented ∼ 40% of the total OL population in the ME at P60+40 (**Figure 1C-D**) compared to just ∼9% of the total OL population in the CC (**Supplementary Figure 1F**). Collectively, these data indicate that new OLs are rapidly and continuously produced in the adult ME.

### 3.2. Oligodendrocytes turn over in the adult median eminence

Despite continuous new OL production, histological assessments in both *Pdgfrα-Cre/ER^T2^;Rosa26-YFP* mice (**Figure 1E**) and a separate cohort of C57BL/6J mice (**Supplementary Figure 1G-I**) indicate that OL numbers are stable in the ME between P60 and P140. Intriguingly, the proportion of YFP-labelled adult-born OLs declines after P60+85 in the ME (**Figure 1D**), but not CC (**Supplementary Figure 1F**). This formed a rationale to directly characterise the stability of ME OLs using *Plp1-Cre/ER^T2^;Rosa26-GFP* mice, in which myelinating OLs are labelled with green fluorescent protein (GFP) following tamoxifen administration (**Figure 1F**). In the ME, the proportion of GFP-labelled OLs present in the ME at the time of tamoxifen administration significantly decreased between P60+9 and P60+37 and remained stable thereafter (**Figure 1G**) despite no differences in the total number of OLs over time (**Figure 1H**). Thus, myelinating OLs are short-lived and continuously replaced in the adult ME. In the CC over the same period, no decline in the proportion of GFP-labelled OLs was observed, as previously reported (*12*), (**Supplementary Figure 1J**). Consistent results were obtained with *Opalin-iCre/ER^T2^;Ai9* mice (alternative OL-specific Cre-driver; **Supplementary Figure 1K-O**), collectively supporting the conclusion that ME OLs turn over in adulthood.

### 3.3. Myelin is continuously replaced in the healthy adult median eminence

We next asked whether ME adult-born OLs become functionally myelinating using *Pdgfrα-Cre/ER^T2^;tau-mGFP* mice, in which tamoxifen administration at P60 induces the expression of membrane-targeted GFP (mGFP) upon OPC differentiation, resulting in the labelling of adult-generated myelin (*22*). Tissues were immunolabelled for mGFP and myelin basic protein (MBP) to distinguish between adult-generated (mGFP^+^/MBP^+^) and pre-existing (mGFP^-^/MBP^+^) myelin (**Figure 2A**). mGFP-labelled myelin rapidly populated the ME, indicating that high rates of local OL production are associated with new myelin formation (**Figure 2B**), and appeared to be specific to the ME since scarce mGFP immunolabelling was detected in other hypothalamic nuclei including the dorso-medial hypothalamus (DMH), ventro-medial hypothalamus (VMH), lateral hypothalamus (LH) and ARC (**Figure 2D**). At P60+80, ∼80% of ME myelin was adult-generated, but this proportion declined thereafter (**Figure 2B**) despite total myelin amounts remaining stable over time (**Figure 2C**). This suggests that in the ME, adult-generated myelin is eventually replaced by new myelin. To test this, we labelled pre-existing myelin at P60 using *Plp1-Cre/ER^T2^;tau-mGFP* mice (**Figure 2E**) and followed its fate over time. In the ME, the amount of mGFP-labelled pre-existing myelin decreased by ∼65% while total myelin amounts remained stable (**Figure 2F-G**). By comparison, previous reports demonstrate that while new myelin is generated after P60 in the CC this occurs at a slower rate and without myelin turnover (*2, 12*). Further supporting local myelin turnover, we observed degraded myelin (dMBP) (*23*) immunolabelling interspersed within the myelin band in the adult ME (**Figure 2H**), indicating myelin degradation. Active phagocytes (Iba1^+^/CD68^+^) as well as myelin-phagocytosing foamy macrophages are also present in the adult healthy ME (**Figure 2I-J**) (*24*). Finally, consistent with data indicating ongoing myelin turnover in the healthy adult ME, we observed MBP immunolabelling inside Iba1+ cells in the ME of healthy adult mice (**Figure 2K-L**, **Supplementary Video 1**), suggesting active phagocytosis of myelin debris by local microglia.

**Figure 2.**
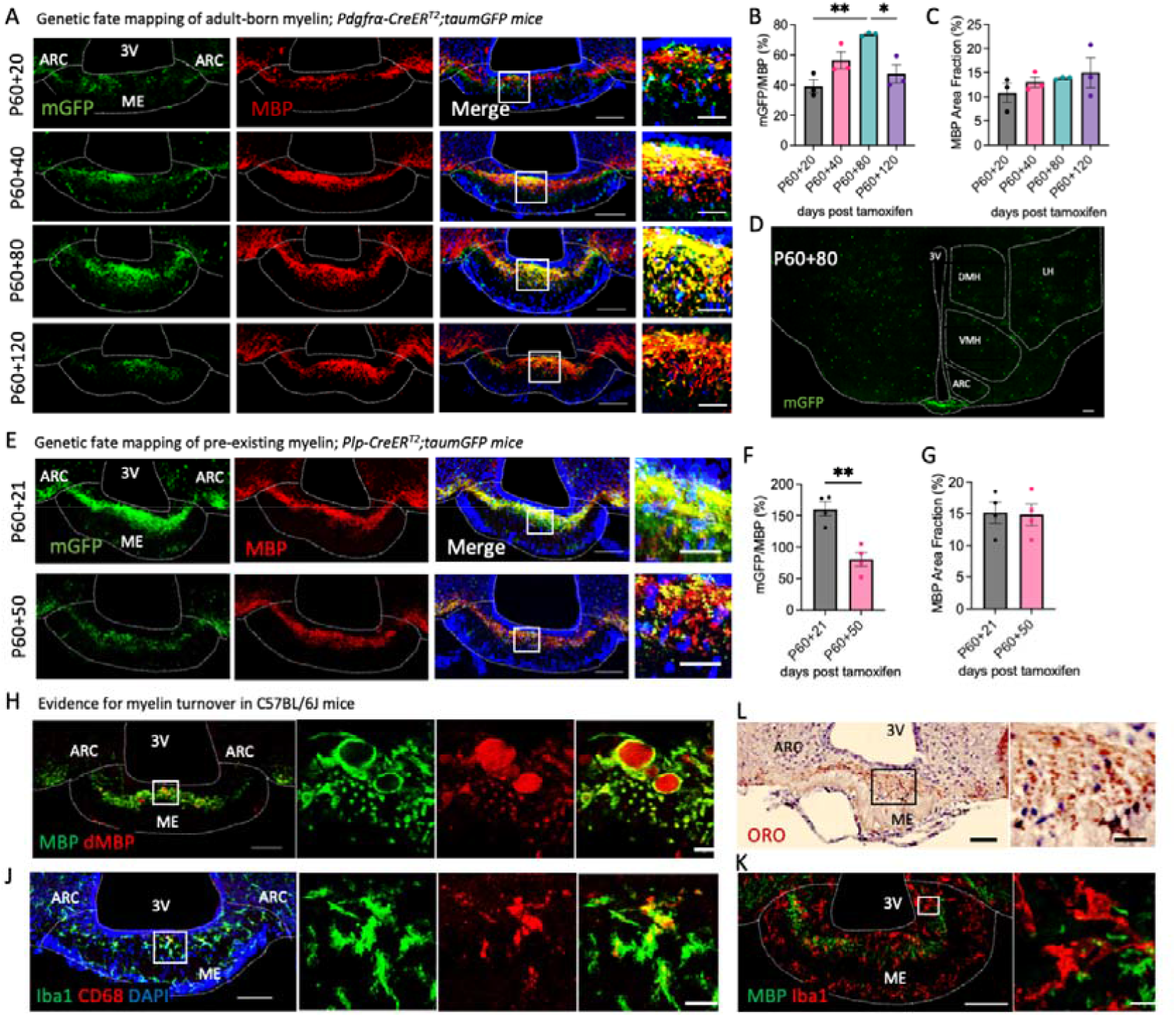
Myelin is continuously generated and replaced in the adult median eminence. Representative images of sections from **(A)** *Pdgfrα-CreER^T2^;taumGFP* and **(E)** *Plp-CreER^T2^;taumGFP* mice immunolabelled for MBP and mGFP and associated quantifications **(B, C, F, G)**. Data (mean ± SEM) were analysed by student’s t-test or one-way ANOVA with Tukey’s multiple comparisons test, *p<0.05, **p<0.01, n=3-4/group. **(D)** shows an overview of mGFP signal in the hypothalamus of a *Pdgfrα-CreER^T2^;taumGFP* mouse at P60+80 (*NB. OPC cell bodies are labelled with a YFP reporter*). Images of ME sections from C57BL/6J adult mice immunolabelled with antibodies against MBP and degraded MBP (dMBP; **H**), oil red o (ORO; **I**) or antibodies recognising Iba1 and CD68 **(J)** or Iba1 and MBP **(K)**. For all images, overview scale bars = 100 μm, inset scale bars = 10 or 20 μm.

### 3.4. Myelin turnover recruits microglia to the adult median eminence

In demyelinating pathologies, myelin debris clearance by phagocytic microglia is critical for new OL formation and myelin repair (*25*). Consequently, we hypothesised that if myelin turnover in the ME requires myelin debris removal by local phagocytes, ablation of microglia would blunt OL and myelin plasticity in the ME. To test this, C57BL/6J mice were treated with PLX5622, a colony stimulating factor 1 receptor inhibitor, to ablate microglia (**Figure 3A-B**). Two weeks PLX5622 treatment reduced MBP immunolabelling in the ME by ∼30% (**Figure 3C-D**), suggesting that local microglia are required for ME myelin maintenance. This was associated with a decrease in OPC proliferation and differentiation, and a reduction in ME OL density (**Figure 3E-F**). Thus, consistent with a role for local microglia in myelin turnover in healthy mice, microglia ablation decreases new OL generation in the ME.

**Figure 3.**
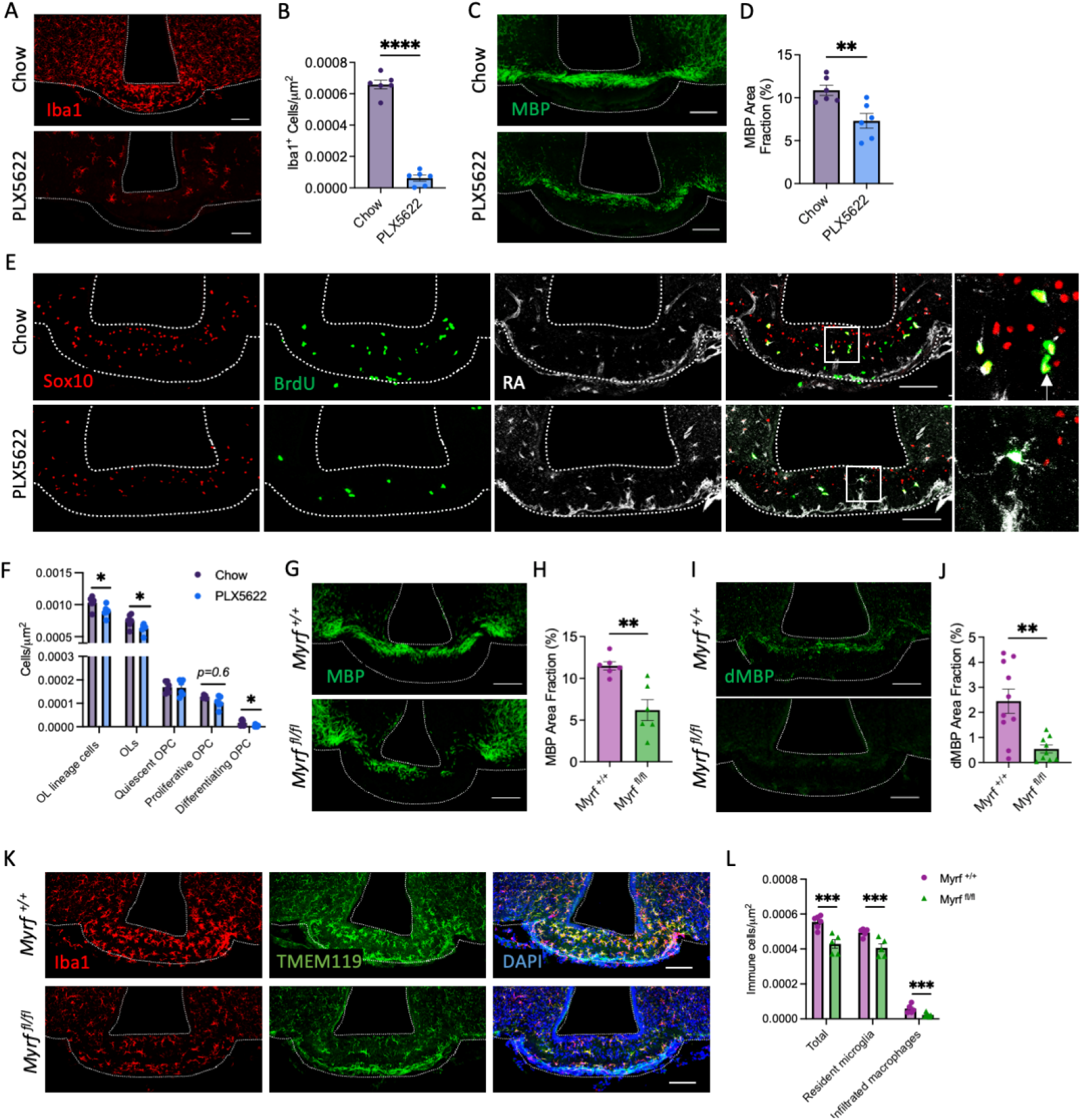
Microglia regulate median eminence oligodendrocyte and myelin plasticity. Representative images of ME sections from C57BL/6J mice fed standard chow or chow containing PLX5622 (12,000 ppm) immunolabelled with antibodies against **(A)** Iba1, **(C)** MBP or **(E)** OL markers and BrdU and associated quantifications **(B, D, F)**. Arrow in **(E)** indicates a differentiating OPC (Sox10+/PDGFRα-/BrdU+). Representative images of ME sections from myelin regulatory factor (*Myrf*) knockout mice (*Myrf ^fl/fl^*) and littermate controls (*Myrf ^+/+^*) immunolabelled with antibodies against **(G)** MBP, **(H)** dMBP and **(K)** Iba1 and TMEM119 and associated quantifications **(H, J, L)**. For all images, overview scale bars = 100 μm, inset scale bars = 20 μm. Data (mean ± SEM) were analysed by student’s t-test or Welch’s test, *p<0.05, **p,0.01, ***p<0.001, n=6-10/group.

Infiltrated macrophages densely populate the ME in healthy mice (*26*), leading us to ask whether myelin turnover and the associated production of myelin debris is responsible for their recruitment. To test this, we used *Pdgfra-Cre/ER^T2^; R26R-eYFP; Myrf^fl/fl^* mice (*Myrf ^fl/fl^* mice), where conditional deletion of myelin regulatory factor (*Myrf*), a transcription factor required for OPC terminal differentiation, blocks new OL formation in the adult brain (**Supplementary Figure 2**). In the ME, *Myrf* deletion rapidly reduced myelin density (**Figure 3G-H**), and myelin debris accumulation (**Figure 3I-J**) and was associated with a decrease in the density of both resident microglia (Iba1^+^/Tmem119^+^) and infiltrated macrophages(Iba1^+^/Tmem119^-^ **Figure 3K-L**). Collectively these data indicate that 1) microglia are required for ongoing ME myelin plasticity and 2) new OL formation is necessary for the the maintenance of the local immune cell population.

### 3.5. Energy availability regulates median eminence myelation

Nutritional signals regulate ME OPC differentiation (Kohnke et al., 2021), but the long-term consequences on ME myelination are unknown. Here we tested whether energy excess or deficit produce changes in ME myelination.

We first assessed the consequences of chronic energy excess on ME myelination using a model of diet-induced obesity (DIO) (**Figure 4A**, **Supplementary Figure 3A**). DIO increased ME OL density and MBP Immunolabelling (**Figure 4B-C**) but paradoxically decreased OPC proliferation and differentiation (**Figure 4D-E**), suggesting that DIO blunts adult oligodendrogenesis in the ME. To test whether increased ME myelination might be the result of changes in OL turnover, we exposed *Plp1-Cre/ER^T2^; R26R-GFP* mice to the DIO paradigm. DIO significantly increased the proportion of YFP-labelled OLs (**Figure 4F-G**) indicating reduced rate of OL turnover.

**Figure 4.**
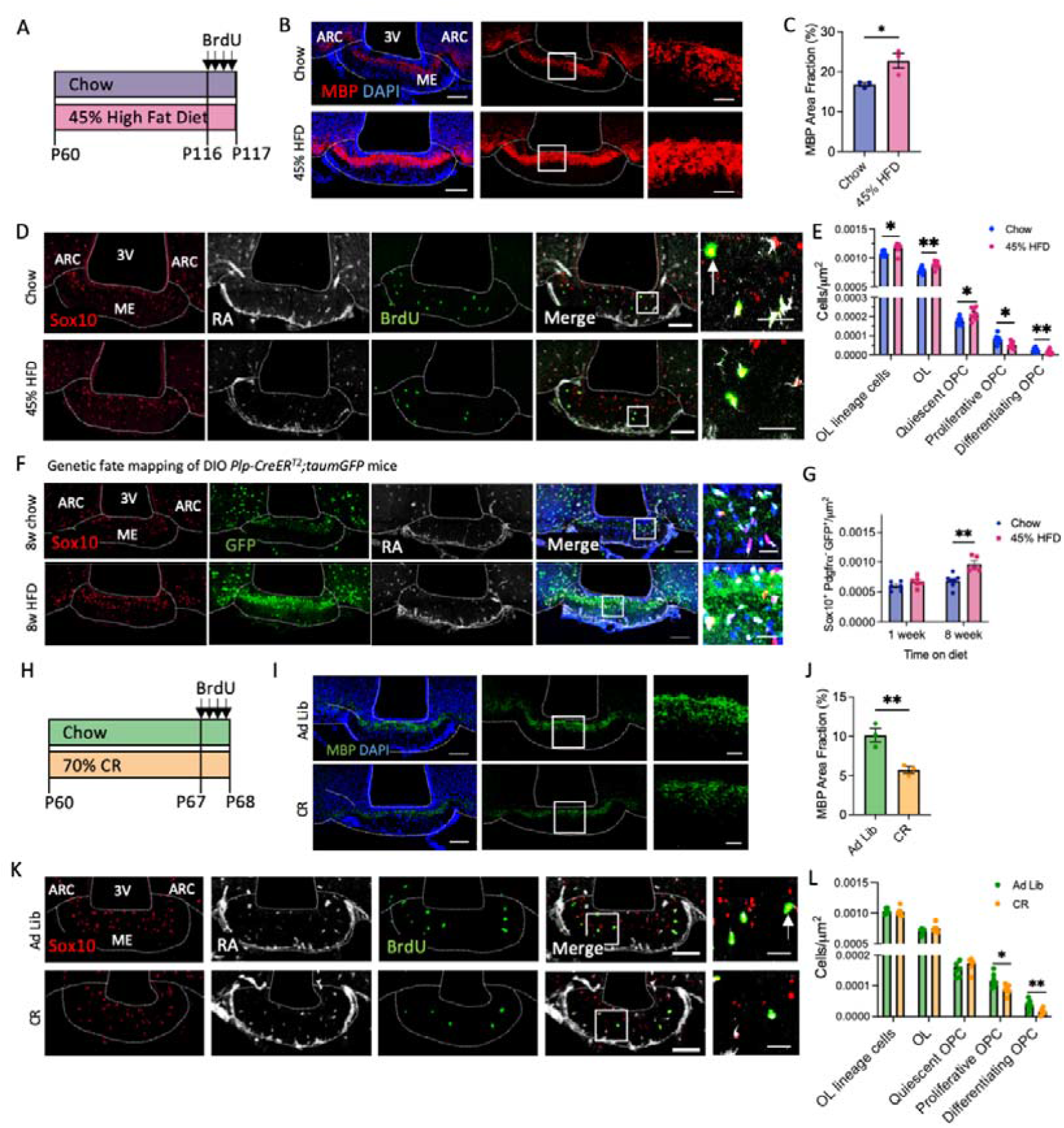
Nutritional regulation of median eminence myelination. Representative images of ME sections from C57BL/6J mice fed a 45% high fat diet (HFD) for 8 weeks **(A)** immunolabelled with antibodies against **(B)** MBP and **(D)** OL markers and BrdU, and associated quantifications **(C, E)**. **(F)** Images of the ME from chow- and 45% *HFD- Plp-CreER^T2^;taumGFP* mice immunolabelled for GFP and OL markers and associated quantification **(G)**. Representative images of ME sections from chow-fed or 70% calorie restricted **(CR, H)** C57BL/6J mice immunolabelled with antibodies against **(I)** MBP and **(K)** OL markers and BrdU and associated quantifications **(J, L)**. For all images, overview scale bars = 100 μm, inset scale bars = 10 or 20 μm. Data (mean ± SEM) were analysed by student’s t-test or two-way ANOVA with Sidak’s multiple comparisons test, *p<0.05, **p,0.01, n=3-6/group.

Finally, we assessed ME myelination in mice exposed to a 7-day 70% calorie restriction (CR) paradigm (**Figure 4H**, **Supplementary Figure 3B**). CR reduced myelin density (**Figure 4I-J**), as well as OPC proliferation (Sox10^+^/PDGFRα ^+^/BrdU^+^) and differentiation (Sox10 /PDGFRα/BrdU) in the ME, resulting in a decrease in OL numbers (Sox10^+^/PDGFRα ^-^, **Figure 4K-L**). Thus, energy deficit reduces new OL production in the ME, resulting in local hypomyelination. Collectively, these data demonstrate that energy availability regulates ME myelination.

## 4. DISCUSSION

Adult oligodendrogenesis is emerging as a new form of brain plasticity observed in a variety of physiological contexts (*8, 10, 11*). In the adult ME, oligodendrogenesis seems to happen within a rapid timeframe with one eighth of OLs being formed within the previous 60 hours, which is unlike what occurs in other brain regions such as the ARC and CC where no adult born OLs are detected within the same period. This high rate of new OL production in the adult ME would have little significance should these new OLs not survive and fully differentiate into myelinating OLs. Six weeks after inducing the labelling of OPCs with YFP, ∼40% of ME OLs are adult-born OLs (**Figure 1**), suggesting that a large fraction of ME adult-born OLs survive. Remarkably, the high rate of new OL formation in the healthy adult ME is offset by a comparably high turnover rate with approximately 40% of ME OLs labelled at P60 disappearing over 6 weeks (**Figure 2**), which sharply contrasts with previous reports of OL longevity in the adult brain (*12*).

How ME OL death is orchestrated and regulated remains to be determined but given the remarkable stability of the number of OLs in the ME, we propose that myelinating OLs may produce a homeostatic feedback signal that regulates both new OL formation and survival, allowing tight control of the OL population. Depletion of microglia impairs this homeostatic control, suggesting that they might form a necessary component of the process. In later adulthood, this homeostatic control seems to be dysregulated (starting P60+120 ± 6 weeks of age), with increased numbers of OLs in the ME, but whether this is due to decreased rates of turnover or increased rates of OL formation remains to be determined. The differentiation potential of OPC might decrease with age in the ME, as has been described in the cortex and CC (*2, 27*). Alternatively, changes in microglial phenotypes with aging (*28*) might also contribute. Interestingly, high fat feeding, which rapidly induces inflammation in this area of the hypothalamus (*29*), blunts OL turnover and the homeostatic control of the ME OL population. Thus, the pro-inflammatory phenotype of hypothalamic microglia might reduce their ability to contribute to OL turnover.

Further studies are needed to elucidate the causes and functional significance of local myelin turnover. Like in other circumventricular organs, the environment of the ME exposes resident cells to lipo- and gluco- toxic insults that might rapidly affect the composition and structure of local myelin, perhaps forming a trigger for OL apoptosis and turnover. However, the observation that nutrient availability regulates ME myelin plasticity suggests this might form a mechanism through which the hypothalamus adapts to changes in energy levels. In fact, previous work identified a role for OPCs in hypothalamic leptin sensing and the regulation of energy balance (Djogo et al., 2016). However, whether the obesity phenotype observed in these studies in response to OPC ablation is induced in response to the loss of OPCs or the subsequent loss of myelin plasticity remains to be clarified.

## Supporting information

Sippl Video 1

## ACKNOWLEDGEMENTS

For the purpose of open access, the author has applied a Creative Commons Attribution (CC BY) licence to any Author Accepted Manuscript version arising from this submission.

We thank the histopathology and imaging cores at the Wellcome-MRC Institute of Metabolic Science for their contributions towards this work. We thank Professors Thora Karadottir and Robin Franklin for useful insights and discussions that have been essential to this project. This work was supported by a Medical Research Council grant (MR/S011552/1; CB), a Wellcome Trust PhD Studentship (108926/Z/15/Z; SB), Medical Research Council Metabolic Disease Unit Grants (MC_UU_00014/5) and (MRC_MC_UU_12012/5), a Wellcome Trust Strategic Award (208363/Z/17/Z) and a Wellcome Trust Investigator Award (108726/Z/15/Z; WDR).

## DATA AVAILABILITY

All data are available in the main text or supplementary materials. Further information and requests for resources and reagents should be directed to and will be fulfilled by the corresponding author, Clemence Blouet.

## AUTHOR CONTRIBUTIONS

Conceptualization: SB, SK, RH, TS, WDR, CBs

Methodology: SB, SK, RH, CB

Investigation: SB, RH, CB

Analysis: SB, CB

Visualization: SB, RH, CB

Funding acquisition: SB, WDR, CB

Writing – original draft: SB, CB

Writing – review & editing: SB, CB

## SUPPLEMENTARY FIGURES

**Supplementary Figure 1.**
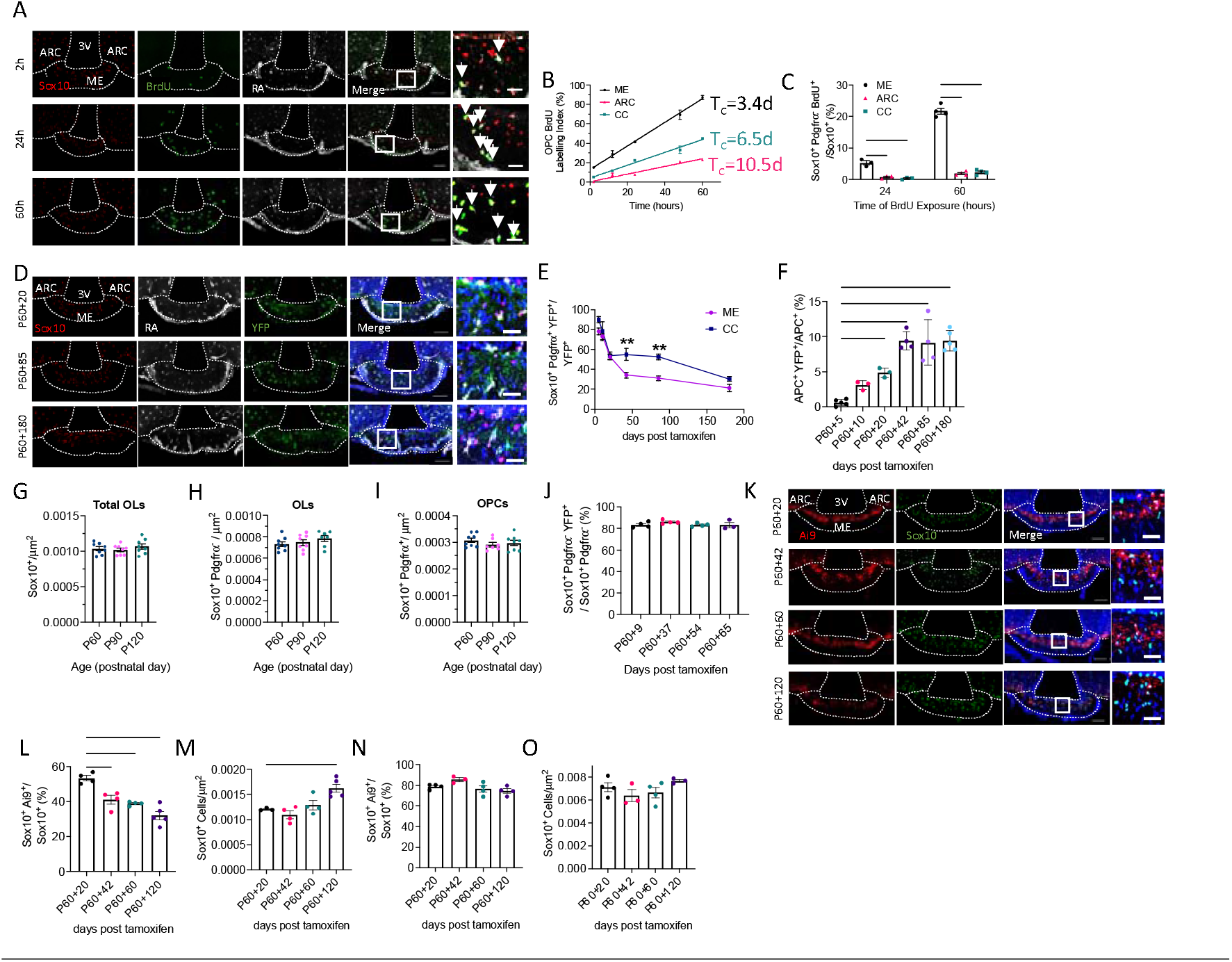
Rapid generation and turnover of oligodendrocytes in the healthy adult median eminence. (**A**) Representative images of OL markers and BrdU in the ME of C57BL/6J mice treated with BrdU. (**B**) The proportion of proliferating OPCs over time was plotted for each region and the resulting graph used to calculate T_C_. (**C**) Quantification of OL lineage cells expressing BrdU but lacking OPC marker expression in the ME, ARC and CC of BrdU-treated C57BL/6J mice. (**D**) Representative images of the ME of *Pdgfra-Cre/ER*^*T2*^*;R26R-YFP* mice immunolabelled for YFP and OL markers and (**E**) the associated quantification of OPC marker expression in YFP+ cells over time. (**F**) Quantification of the proportion of adult born OLs generated from OPCs after P60 in the CC of *Pdgfra-Cre/ER*^*T2*^*;R26R-YFP* mice. Quantification of (**G**) OL lineage cells, (**H**) OLs and (**I**) OPCs in the ME of C57BL/6J mice at P60, P90 and P120. (**J**) Quantification of YFP-labelled OLs in the CC of *Plp-Cre/ER*^*T2*^*;R26R-eG*FP –mice following tamoxifen administration at P60. (**K**) Representative images of Ai9 expression in OLs in the ME of *Opalin-CreER*^*T2*^*;Ai9* mice following tamoxifen administration and associated quantifications (**L**,**M**). Quantification of (**N**) Ai9 reporter expression in OLs and (**O**) OLs in the CC of *Opalin-CreER*^*T2*^*;Ai9* mice. For all images, overview scale bars = 100 μm, inset scale bars = 20 μm. Data analysed by one-or two-way ANOVA with post-hoc analysis by Dunnett’s or Sidak’s multiple comparisons test, **p,0.01, ***p<0.001, ****p<0.0001, n=3-8/group.

**Supplementary Figure 2.**
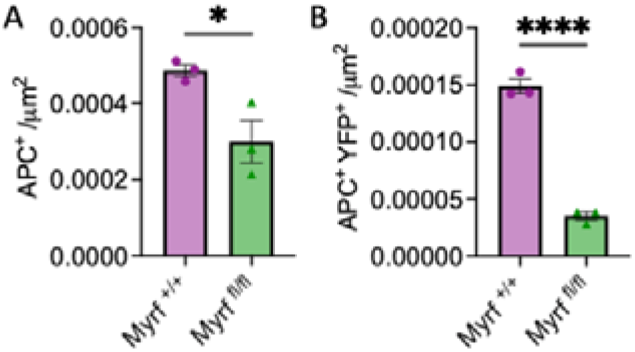
Adult-onset deletion of myelin regulatory factor blunts oligodendrogenesis in the adult median eminence. Quantification of the density of **(A)** mature oligodendrocytes expressing APC and **(B)** adult-born oligodendrocytes in the median eminence of myelin regulatory factor knockout (*Myrf ^fl/fl^*) mice and wild-type controls (*Myrf ^+/+^*) three weeks after tamoxifen administration at P60. Analysed by student’s t-test, *p<0.05, ****p<0.0001, n=3/group.

**Supplementary Figure 3.**
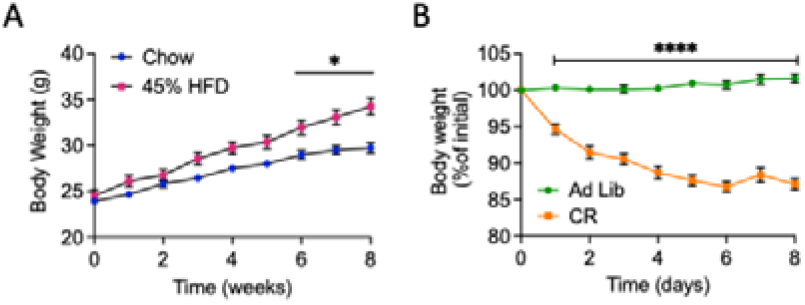
Effects of nutritional interventions on body weight. Body weight of C57BL/6J mice fed a **(A)** control or 45% high fat diet (HFD) for 8 weeks or **(B)** chow diet ad libitum (AL) or 70% calorie restricted (CR) for 7 days. Data analysed by repeated measures two-way ANOVA, *p<0.05, ****p<0.0001, n=8/group.

**Supplementary Video 1:** Engulfment of myelin by median eminence microglia. Video demonstrating MBP immunolabelling (green) inside Iba1^+^ microglia (red) in the median eminence of healthy, chow-fed animals.

